# Integrating multimeric threading with high-throughput experiments for structural interactome of *Escherichia coli*

**DOI:** 10.1101/2020.10.17.343962

**Authors:** Weikang Gong, Aysam Guerler, Chengxin Zhang, Elisa Warner, Chunhua Li, Yang Zhang

**Affiliations:** Department of Computational Medicine and Bioinformatics, University of Michigan, Ann Arbor, Michigan, 48109, USA; Faculty of Environmental and Life Sciences, Beijing University of Technology, Beijing 100124, China; Department of Biological Chemistry, University of Michigan, Ann Arbor, Michigan, 48109, USA

**Keywords:** Protein-protein interaction networks, multiple-chain threading, *Escherichia coli* genome, structural interactome, network centrality

## Abstract

Genome-wide protein-protein interaction (PPI) determination remains a significant unsolved problem in structural biology. The difficulty is twofold since high-throughput experiments (HTEs) have often a high false-positive rate in assigning PPIs, and PPI quaternary structures are more difficult to solve than tertiary structures using traditional structural biology techniques. We proposed a uniform pipeline to address both problems, which first recognizes PPIs by combining multi-chain threading alignments with HTE results using naïve Bayesian classifiers, where the quaternary complex structures are then constructed by mapping the monomer models with the dimeric threading frameworks through interface-specific structural alignments. The pipeline was applied to the *Escherichia coli* genome and created 35,125 confident PPIs which is 4.5-fold higher than HTE alone. Graphic analyses of the PPI networks revealed a scale-free cluster size distribution, which was found critical to the robustness of genome evolution and the centrality of functionally important proteins that are essential to *E. coli* survival. Furthermore, complex structure models were constructed for all predicted *E. coli* PPIs based on the quaternary threading alignments, where 6,771 of them were found to have a high confidence score that corresponds to the correct fold of the complexes with a TM-score >0.5 and 93 showed a close consistency with the later released experimental structures with an average TM-score=0.73. These results demonstrated the significant usefulness of threading-based homologous modeling in both genome-wide PPI network detection and complex structural construction.

## INTRODUCTION

Most proteins conduct functions through interactions, either permanently or transiently, with other proteins. These interactions result in various protein-protein interaction (PPI) networks, or interactomes (Huttlin et al. 2015), that are essential to accommodate many important cellular processes, ranging from transcriptional regulation to signal transduction and metabolic pathways. Experimental methods to elucidate these networks are, however, limited and most of them (including yeast-two hybrid and tandem-affinity purification) have high error rates up to 90% (von Mering et al. 2002). Furthermore, these high-throughput experimental (HTE) methods only address the issue of what proteins interact, but cannot provide information as to where and how the proteins interact; this information is critical for understanding the biophysical mechanisms of the interaction networks and/or developing new therapies to regulate the networks (Archakov et al. 2003).

While structure biology through X-ray and NMR techniques could in principle provide the most accurate structural information of PPIs, these experiments are however often too expensive and labor intensive to be applied on a genomic scale. There are also many complexes that are currently difficult to solve due to technical difficulties in protein expression and crystallization. In *Escherichia coli*, the most studied bacterial organism of our time, for example, there are only 1,450 out of the 4,280 protein-coding genes (<34%) that have the structures experimentally solved (Xu and Zhang 2013). The number of PPI complex structures is even less, with 693 PPI entries in the PDB which counts only for <7% of the ~10,000 putative PPIs in *E. coli* (Rajagopala et al. 2014). Homology modeling has been proved to be an effective approach to construct structure models by copying the frameworks from homologous PPI templates (Szilagyi and Zhang 2014). But until recently, the approach did not significantly contribute to the elucidation of PPI networks, due to the limited number of available homologous complex structures in the PDB (Aloy et al. 2004; Kundrotas et al. 2012; Szilagyi and Zhang 2014). Recent studies have shown that the structural library of PPI interfaces approaches to completion (Gao and Skolnick 2010), implicating that most of the complexes should have analogous interfaces in the PDB; this settles a promising base for the template-based structure modeling of a wide-range of interactions if advanced threading methods can be developed to recognize such analogies. There are also excellent efforts that tried to combine interaction data from different resources for large-scale PPI network identification (Jansen et al. 2003; von Mering et al. 2005; Zhang et al. 2012); many of the approaches however do not provide 3D structures of the complexes.

In this work, we proposed a new hybrid pipeline, Threpp, which extends the multiple-chain threading protocol (Guerler et al. 2013) to address two central problems of protein interactomes (Figure 1). First, we will develop a new Bayes classifier model to integrate high-throughput proteomic data with multimeric threading alignments to improve the accuracy and coverage of PPI recognitions. 3D structures of protein complexes are then constructed for all the predicted PPI pairs by threading the query sequences through a non-redundant complex structure library. Different from several existing homology-based methods that build complex structures by multiple-chain sequence comparison (Lu et al. 2002; Mukherjee and Zhang 2011), which requires separate complex library construction and often misses specific binding modes, Threpp deduces complex structure templates directly from monomer chain threading followed by oligomer-based mapping, which enables the multiple binding mode recognition through the entire PDB library. It is also different from the template-based docking (Kundrotas et al. 2012; Zhang et al. 2012; Szilagyi and Zhang 2014) which associates monomer and dimer structures by pure structural similarity, while Threpp detects PPI frameworks and the monomer-dimer associations using profile-based threading alignments which often have a higher accuracy than pure structure comparisons.

**Figure 1.**
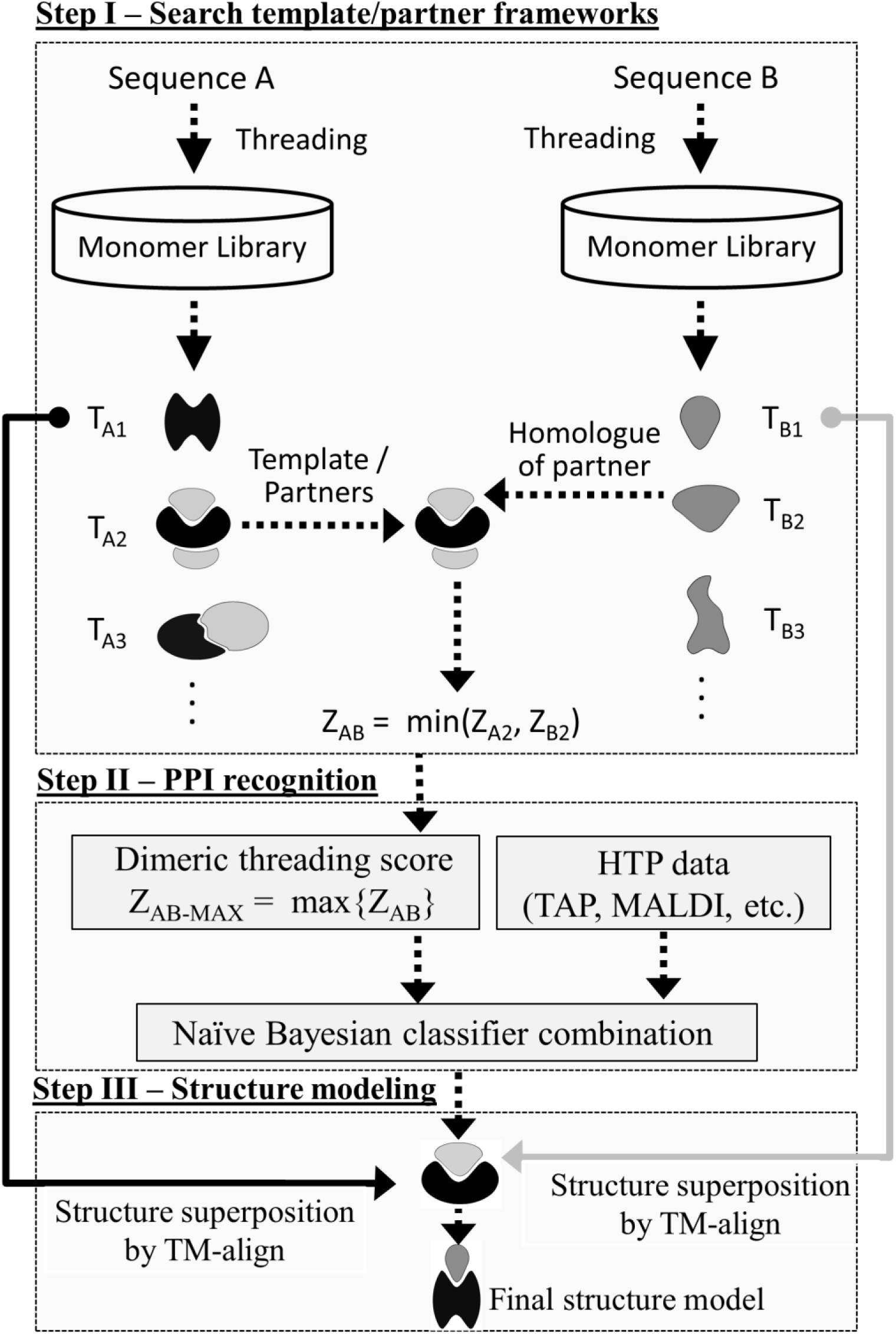
Flowchart of Threpp for PPI recognition and structure construction. The pipeline consists of three steps of threading-based PPI framework identification, Bayesian classifier PPI recognition, and PPI complex structure construction by monomer/dimer template recombination.

To examine the accuracy of Threpp, we carefully benchmarked the strength and weakness of the pipeline in PPI recognition on large-scale gold standard datasets. As a case study, the pipeline was applied to the *E. coli* genome to construct the structural networks of the species, with results revealing important functional implications of the modeled interactome. The Threpp algorithm, together with the structural models of all PPIs for the *E. coli* genome, are made freely downloadable to the community at genome, are made freely downloadable to the community at https://zhanglab.ccmb.med.umich.edu/Threpp/.

## RESULTS

### Benchmark test of Threpp on PPI assignments

To train and test Threpp for PPI recognitions, we collected a ‘Gold Standard’ (GS) set of PPIs in the *E. coli* that have definite positive and negative references as assigned by Hu et al (Hu et al. 2009), where the positive samples contain 763 experimentally-established physical interactions obtained from DIP (Xenarios et al. 2000), BIND (Bader et al. 2003) and INTACT (Kerrien et al. 2012) databases, and the negative set consists of 134,632 putatively non-interacting protein pairs compiled from the protein pairs belonging to different cellular compartments (see Table S1 in Supplementary Information, SI). Here, membrane proteins were excluded due to the close physical proximity (and potential physical interaction) with both cytoplasmic and periplasmic proteins.

#### PPI recognition by individual threading and HTE methods

Figure 2 presents the true positive rate (TPR) and false positive rate (FPR) of PPI assignments for the test proteins by Threpp based only on the Z-score of dimeric threading alignments, *Z*_*com*_ (named as ‘Threpp_threading’, see Methods), where the detail of the data is listed in Table S2. Here, 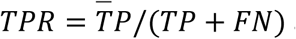 and *FPR* = *FP*/(*FP* + *TN*), with the standard true positive (*TP*), true negative (*TN*), false positive (*FP*) and false negative (*FN*) calculated by comparing the PPI predictions with the GS assignments. As a comparison, we also list in Figure 2 the results from four sets of HTEs, including two tandem-affinity purification (TAP) sets (‘Butland set’ (Butland et al. 2005) and ‘Hu set’ (Hu et al. 2009)), the ‘Arifuzzaman set’ derived through matrix-assisted laser desorption/ionization time-of-fight (MALDI-TOF) mass spectrometry (Arifuzzaman et al. 2006), and the ‘Rajagopala set’ obtained by yeast two-hybrid (Y2H) screening (Rajagopala et al. 2014), where detailed assignments by the HTE data are listed in Table S3.

**Figure 2.**
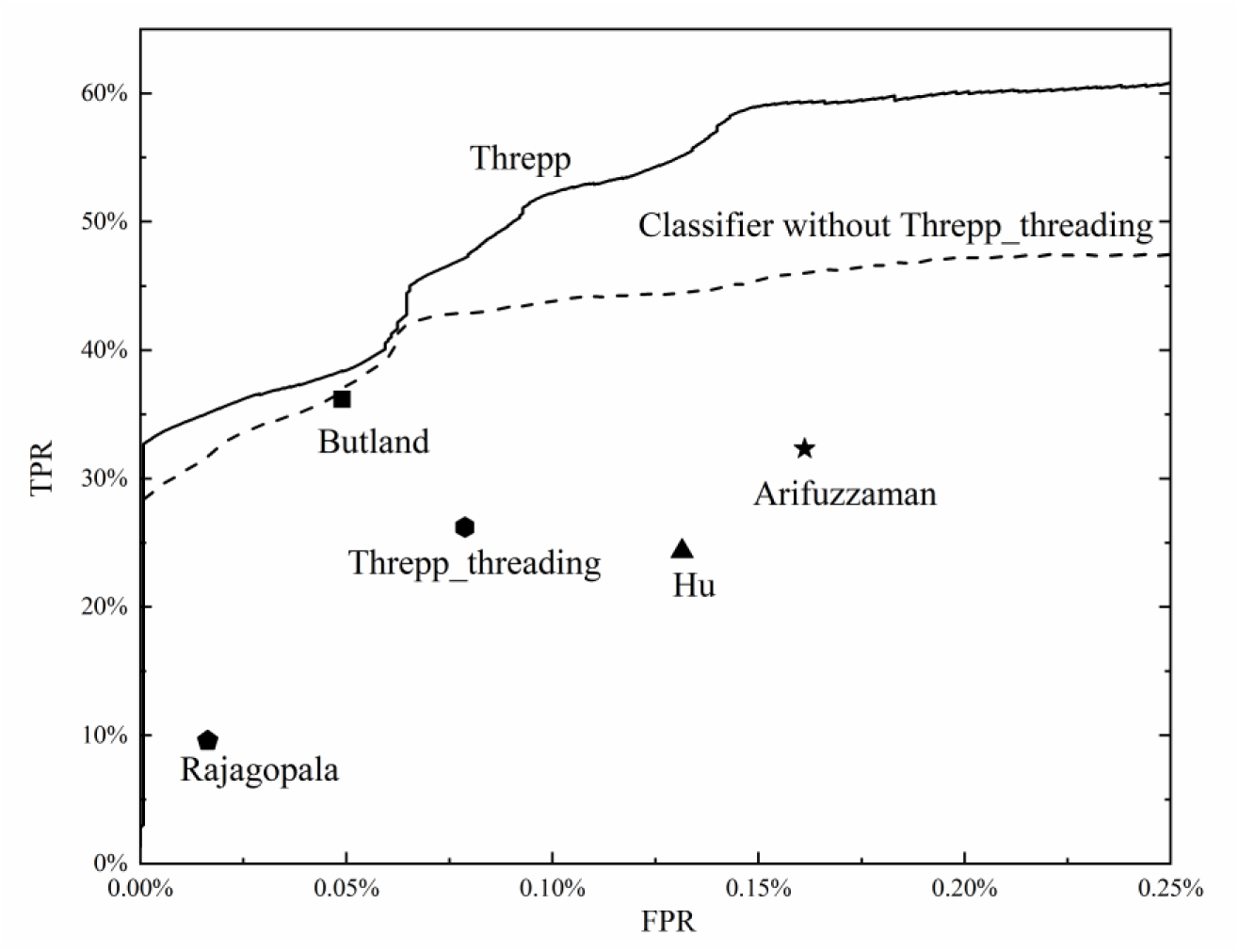
True positive (TPR) and false positive (FPR) rates of PPI recognition by different approaches. The predictions of Bayesian classifiers combining different sources of interaction evidences, with and without Threpp_threading, are shown in solid and dashed lines, respectively.

Table 1 (upper panel) summarizes the Matthew’s correlation coefficient (MCC) by the individual methods, where 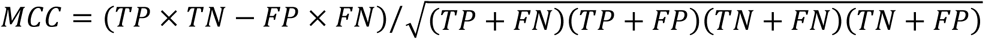 represents a balanced metric of precision and recall of the PPI predictions. While the MCC of Threpp_threading (0.41) is lower than that of the ‘Butland set’ (0.54), it is comparable or slightly higher than other HTE results, including the ’Arifuzzaman set’ (0.41), the ’Hu set’ (0.35), and the ‘Rajagopala set’ (0.27).

**Table 1.** Summary of PPI recognition by different methods.

|  | MCC | TPR | FPR |
| --- | --- | --- | --- |
| *Individual datasets from high-throughput experiments and threading* |  |  |  |
| Tandem affinity purification (Butland set) | 0.54 | 36.2% | 0.05% |
| MALDI-TOF (Arifuzzaman set) | 0.41 | 32.4% | 0.16% |
| Tandem affinity purification (Hu set) | 0.35 | 24.4% | 0.13% |
| Yeast-two hybrid (Rajagopala set) | 0.27 | 9.6% | 0.02% |
| Threpp_threading | 0.41 | 26.2% | 0.08% |
| *Bayes combinations* |  |  |  |
| Classifier without Threpp_threading | 0.58 | 42.4% | 0.07% |
| Threpp | 0.64 | 59.1% | 0.14% |

#### Bayesian classifier models increase PPI recognition accuracy of individual methods

In the lower panel of Table 1, we list the results of combined data from different methods by Threpp. First, we used the Bayesian classifier to combine the data from 4 HTE datasets, which results in a higher MCC (=0.58) than all individual datasets. As shown in Figure 2 (the dashed curve), both TPR and FPR increase with the decrease of the threshold, but the curve is above all individual experimental datasets, demonstrating the effectiveness of the Bayesian classifier model in selecting correct PPIs. Nevertheless, the MCC difference between the Bayesian model (0.58) and the best HTE data from the ‘Butland set’ (0.54) is modest.

After combining the HTE with the Threpp_threading models, the MCC is increased to 0.64, which is 18.5% higher than the best individual dataset from the ‘Butland set’. This difference in MCC corresponds to a p-value = 1.1E-59 in the Student’s t-test, indicating that the difference is statistically significant. The result suggests that although the accuracy of the threading score is not high on its own, the modeling data is highly complementary to the HTE evidences, where a naïve Bayesian combination of the computer-based and experimental data can thus result in a highly significant improvement of both the recall and the precision of the PPI predictions.

### Integrating threading model with HTE data for *E. coli* network detection

The *Escherichia coli* genome contains in total 4,280 protein-coding genes (Benson et al. 2018). As an application, we used Threpp to evaluate all the 9,157,060 putative pairs by the Bayesian combination of dimeric threading and HTE datasets (Butland et al. 2005; Arifuzzaman et al. 2006; Hu et al. 2009; Rajagopala et al. 2014). Despite the huge number of putative interactions, only 4,280×2 monomer threading runs are needed with the interaction frameworks assigned by a pre-calculated homology look-up table for all templates, where the genome-scale network calculation is fast with ~2 hours on a 2000-HPDL1000h core cluster.

The experiment yielded 35,125 confident PPIs (Table S4), which has a likelihood rate score above 1.87 by Threpp (see Eq. 3 in Methods). In case where the HTE data are not available, only Threpp threading scores are employed for the targets with a stringent complex framework Z-score cutoff of *Z*_*com*_ ≥ 25. Our benchmark results on the GS datasets show that the PPI assignments with such likelihood score and *Z*_*com*_ cutoffs have an average accuracy of 0.996. Overall, these interactions are combined from 28,263 PPIs by Threpp_threading and 21,932 by the four HTE datasets, where there are only 1,153 PPIs in the intersection set of the two and 13,917 were dropped off by Threpp due to insufficient likelihood rate score. These predicted interactions contain 451 out of the 763 PPIs in the GS dataset, which is significantly higher than the number of GS PPIs predicted by either Threpp_threading (200) or the four HTE approaches (346).

Here, if we ignore the threading alignments and only combine the HTE data, the number of PPIs detected by Threpp will be reduced to 7,872 that have the similar level of likelihood score, which corresponds to only 22% of all PPIs identifiable by the full Threpp pipeline. These data demonstrate again a high complementarity of the threading alignments to the HTE data, and in particular the impact of consideration of threading-based approach on the hybrid PPI recognitions.

### PPI networks reveal dominant roles of essential proteins in *E. coli*

The 35,125 high-confidence PPI assignments detected by Threpp involve 3,273 proteins. Based on these PPIs, we constructed a comprehensive *E coli* protein interaction network (Figure 3a). In the plot, nodes represent individual proteins with edges being the interactions between proteins, where self-loops (corresponding to orphan proteins) and multiple edges (repeated PPI predictions) have been excluded.

**Figure 3.**
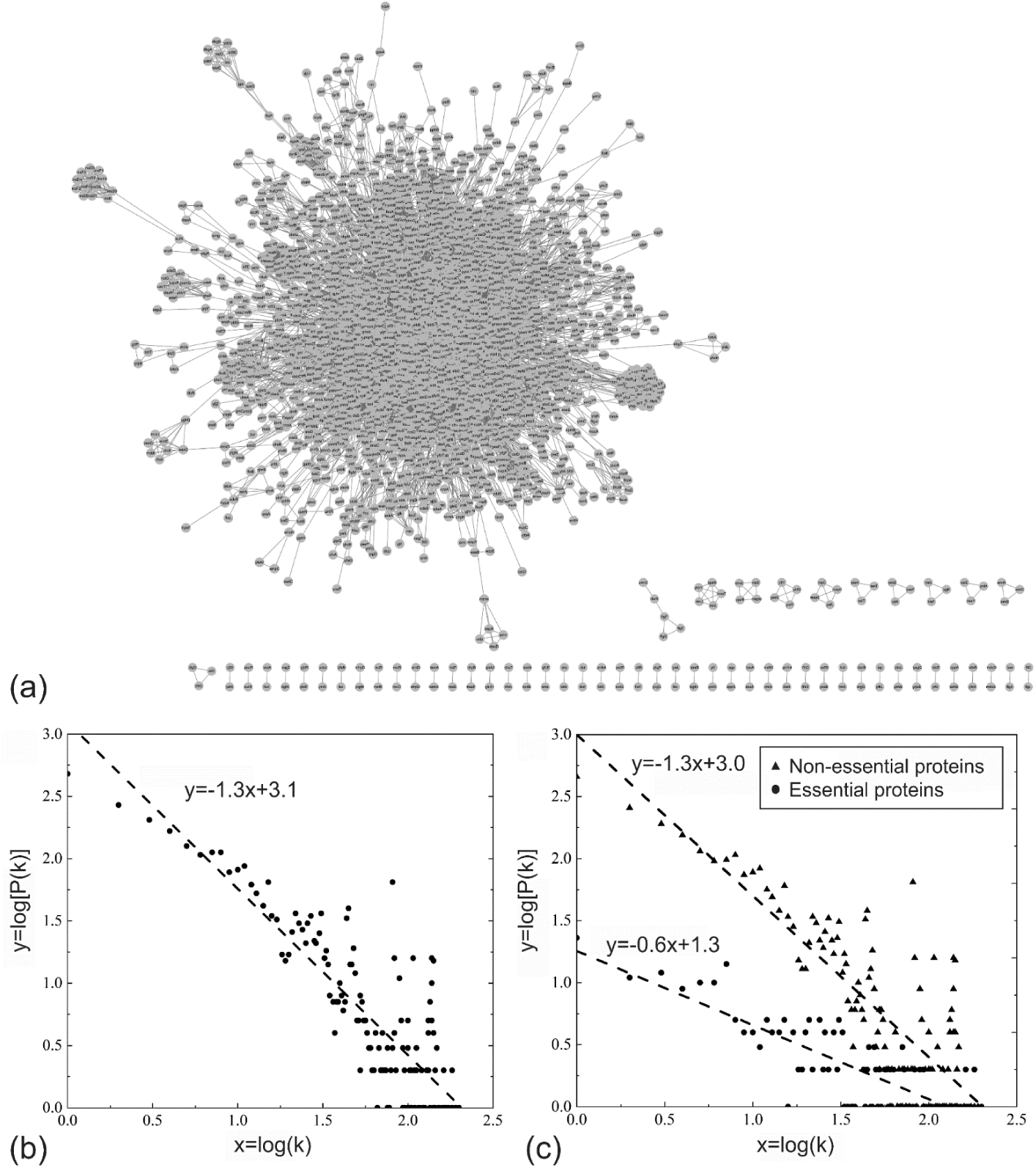
PPI networks and degree distributions of the *E coli* genome. (a) PPI networks constructed from 35,125 high-confidence PPIs by Threpp, which involve 3,273 proteins. (b) Distribution of PPI node degree (*k*) that is defined as the number of edges cross each node in the network. (c) PPI cluster distribution for essential (circles) and non-essential (triangles) proteins. Lines in (b) and (c) are power law fit to *P*(*k*) ∝ *k*^−*γ*^.

#### Node degree distribution is scale free

Figure 3b shows the degree distribution of the PPI networks for all 3,273 involved proteins, which follows a power-law of *P*(*k*) ∝ *k*^−1.33^, where the degree (*k*) of a protein node equals to the number of edges that have this node as one of its endpoints. This network possesses two outstanding characteristics which are important to facilitate the biological functionality and evolution of the *E coli* genome. First, there are dominantly more proteins in the genome with few interaction partners; this property of PPI networks helps enhance the robustness of the network against random mutations in the evolution, as the overall network is not influenced by the deletion and insert of individual proteins. On the other hand, the scale-free nature of the degree distribution indicates that a non-trivial number of proteins, which is significantly higher than what is expected from a normal distribution, have many interaction partners; this feature allows a substantial amount of important proteins to serve as hub of interactions and dominate the functional interaction networks.

#### Essential proteins interact with more partners than non-essential ones

In Figure 3c, we present the degree distributions of PPI networks for two sets of essential and non-essential proteins separately, where the 303 essential proteins are taken from Baba *et al*. that were found unable to be deleted from the chromosome for the survival of *E. coli* through the large-scale gene-deletion assay, and the rest are considered as ‘non-essential’ (Baba et al. 2006). While both protein sets follow a stringent power-law distribution (i.e., *P*(*k*) ∝ *k*^−0.60^ for essential and *P*(*k*) ∝ *k*^−1.34^ for non-essential proteins), the average connectivity (or degree *k*) per node is significantly higher for essential proteins (33.5) than for non-essential genes (18.5). In particular, the percentage of proteins with more than 34 interaction partners in the essential proteins (30%) is much larger than that in the non-essential proteins (15%), indicating that the essential proteins tend to serve as the interaction hub which has resulted in their significant functional importance for *E. coli* to survive.

#### Betweenness centrality

To examine the centrality of proteins in the PPI network, we define the betweenness centrality (BC) of a protein node *ν* by (Freeman 1977):

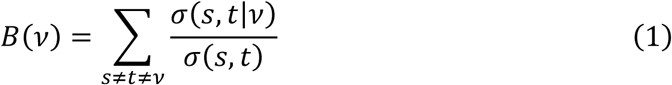

where *σ*(*s*, *t*) denotes the number of shortest paths from nodes *s* to *t*, and *σ*(*s*, *t*|*ν*) is the number of the shortest paths from *s* to *t* that cross through *ν*. The sum in Eq. (1) runs through all node pairs in the network excluding the target node *ν*. Here, although both BC and degree (*k*) defined above are related to the number of interaction partners for a given protein node, BC measures the number of the shortest paths passing through one node and reflects the information flow through the protein, i.e., a protein with a higher BC tends to control more functional flow of the PPI networks.

In Table S5, we list the BC values for all protein nodes in *E coli* that have at least one interaction partner in the Threpp predicted PPI networks. The top ten nodes with the highest BCs are presented in Table 2, which all correspond to the functionally important proteins involving complex cellular processes, including chaperone, elongation factor, transcriptional regulatory and ribosomal proteins.

**Table 2.** The ten proteins with the highest betweenness centrality (BC) values.

| ID | BC | Name of proteins |
| --- | --- | --- |
| DnaK | 0.049 | Chaperone protein DnaK |
| TufA | 0.037 | Elongation factor Tu 1 |
| RpsB | 0.029 | 30S ribosomal protein S2 |
| MetN | 0.029 | Methionine import ATP-binding protein MetN |
| LpdA | 0.027 | Dihydrolipoyl dehydrogenase |
| RplL | 0.027 | 50S ribosomal protein L7/L12 |
| TufB | 0.027 | Elongation factor Tu 2 |
| RlmN | 0.020 | Dual-specificity RNA methyltransferase RlmN |
| RplV | 0.018 | 50S ribosomal protein L22 |
| RcsB | 0.018 | Transcriptional regulatory protein RcsB |

As an illustration, we present in Figure 4 a local PPI network involving the DnaK protein, which has the highest BC score (=0.049). DnaK is known to serve as a chaperone to promote protein folding, interaction and translocation, both constitutively and in response to stress, by binding to unfolded polypeptide segments (Zhu et al. 1996). Here, RcsA, RcsB (with the 10th largest BC value) and RcsD are all involved in the Rcs phosphorelay pathway, a complex signal transduction system. Through this pathway, phosphate travels from the phosphotransfer protein RcsD to RcsB, which is essential to the regulation of a variety of cellular processes in the bacteria. In this example, the BC-based analysis helps to reveal the key role that the DnaK protein exerts in connecting the metabolic pathway (Rcs phosphorelay pathway) and cell process (cell division regulated by gene ftsA) (Carballes et al. 1999). With the PPI network data provided by the Threpp modeling, the BC analysis can be extended to other systems for key protein and pathway identifications to facilitate various medical and pharmaceutical studies.

**Figure 4.**
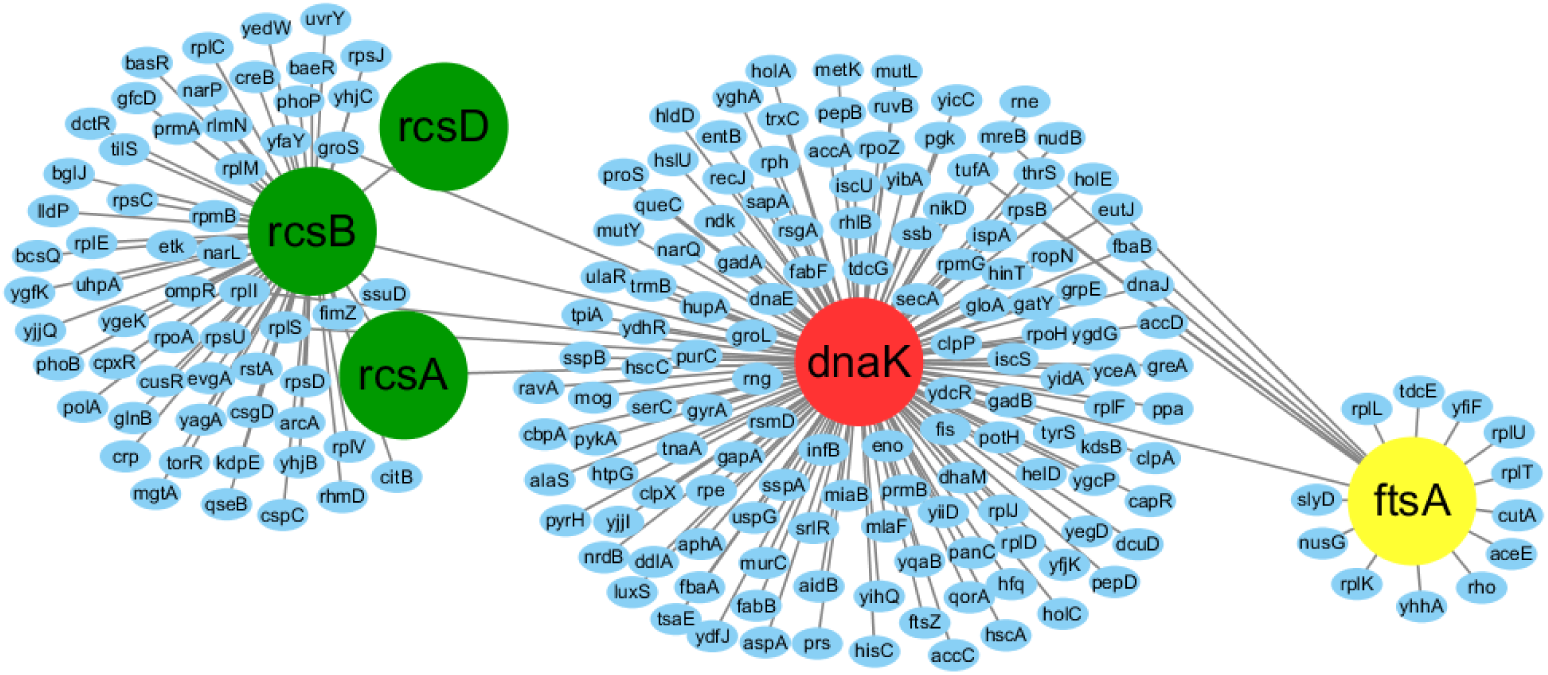
A local PPI network involving the chaperone protein DnaK (red) that mediates the Rcs phosphorelay signaling pathway. The RcsA, RcsB and RcsD proteins (in green) positively regulate the expression of the cell division gene ftsA (yellow) through the interactions with the DnaK protein.

### Structural modeling of protein interactome in *E. coli*

For structural interactome, Threpp was used to create 3D structure models for all the predicted 35,125 PPIs (see http://zhanglab.ccmb.med.umich.edu/Threpp/Ecoli3D.zip), where 6,771 are found to have a Threpp S-score >13 (see https://zhanglab.ccmb.med.umich.edu/Threpp/download/Ecoli3D.txt for S-score values of all the PPI complexes). Here, S-score is defined in Eq. (4) in Methods for estimating model quality of Threpp predictions. In a previous benchmark study (Guerler et al. 2013), it was shown that 78% of the dimer-threading models with a S-score >13 can have a TM-score >0.5 to their co-crystallized reference structures, indicating correct quaternary structure fold (Xu and Zhang 2010). Below, we selected two complexes from a DMSO reductase (DmsAB) and a hetero-trimeric xanthine dehydrogenase (YagRST), as illustrative examples (S-score >13) to analyze in detail the Threpp models. Although these PPIs have been shown critical to the function of *E. coli*, the interactions were not reported by any of the four high-throughput experimental datasets.

#### Dimethyl sulfoxide reductase complex (DmsAB)

*E. coli* is well known to withstand anaerobic conditions through the utilization of correlated reductases in anaerobic media, while DmsAB is a critical dimethyl-sufoxide reductase complex that supports the bacterial growth in anaerobic media via electron transport. Although no structure has been solved for any of the protein components, there are several experimental evidences that can be used as indirect validations of the Threpp structure modeling. For example, DmsA and DmsB are known to contain one (FS0) and four [4Fe-4S] clusters (FS1 to FS4) respectively for electron shuttling (Rothery et al. 2008), and DmsB is anchored on the membrane via residues Pro80, Ser81, Cys102 and Tyr104, where these resides are also used for mediating the downstream electron transferals (Cheng et al. 2005).

Figure 5a shows a cartoon representation of the Threpp model for the DmsAB complex, which has a high-confidence S-score of 52.3. The monomeric structure models for DmsA and DmsB were derived from the templates of PDB ID 1EU1 (chain A) and 2VPZ (chain B), while the orientation of the monomers was modeled using 2IVF (chains A and B) that was recognized by Threpp_threading as dimeric framework (see Table S6). Although the monomer and complex templates have been identified separately, the TM-score and RMSD of the predicted model from the dimer framework are 0.89 and 3.54 Å, respectively, showing a high consistency of the monomer threading and dimeric framework. The framework protein, 2IVF, is a member of the DMSO reductase family and serve as an Ethylbenzene Dehydrogenaes from *Aromatoleum aromaticum*. As highlighted in Figure 5a, the complex model for DmsAB also contains well-shaped five [4Fe-4S] clusters, demonstrating the close consistency with the insights from the biochemical experiments (Cheng et al. 2005; Rothery et al. 2008).

**Figure 5.**
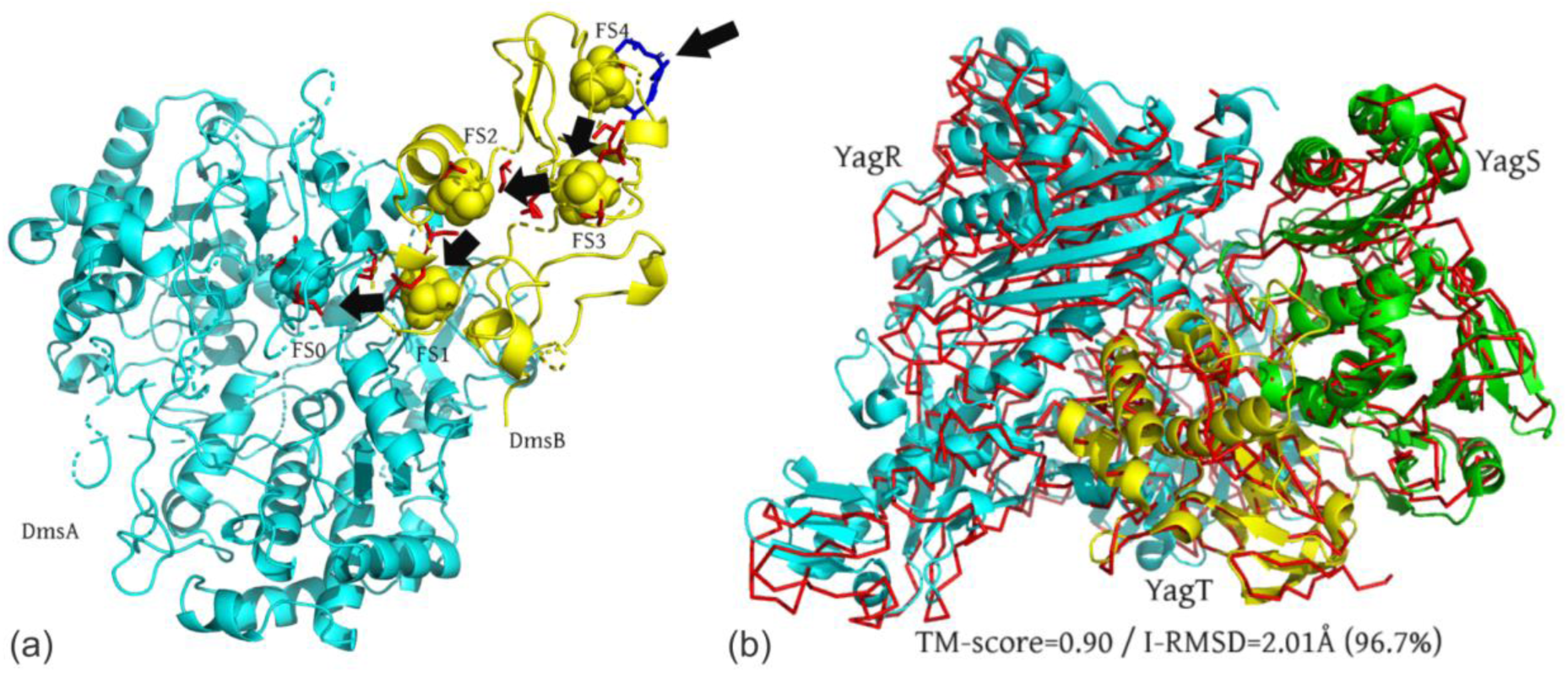
Illustrative examples of quaternary structure models by Threpp. (a) DmsA (cyan) and DmsB (yellow) complex, where the predicted [4Fe-4S] clusters (FS0 to FS4) are highlighted as spheres with arrows indicating the direction of electron transportation. The binding sites of the [4Fe-4S] clusters determined by biochemistry experiments (Rothery et al. 2008) are shown as red sticks, where the membrane anchor residues Pro80, Ser81, Cys102 and Tyr104 in DmsB are shown as blue sticks. (b) Trimeric iron-sulfur complex YagRST, where the Threpp model (red lines) is superposed on the X-ray structure (cartoon) that was solved after the modeling was completed. The monomer chains of YagR, YagS and YagT are shown in blue, green and yellow, respectively.

#### Trimeric iron-sulfur complex (YagRST)

YagRST is a molybdenum-containing iron-sulfur enzyme located in the periplasm of *E. coli*, which functions in cell maintenance by detoxifying aromatic aldehydes to avoid cell damage (Neumann et al. 2009). Structurally, YagRST is a heterotrimer complex consisting of a large 78.1 kDa molybdenum-containing subunit (YagR), a medium 33.9 kDa FAD-containing subunit (YagS), and a small 21.0 kDa 2Fe2S-containing subunit (YagT). Built on the threading alignments, Threpp first created monomeric structure models for YagR, YagS and YagT using templates with PDB ID 1RM6 (chain A), 1RM6 (chain B) and 3SR6 (chain A), respectively. Accordingly, three framework templates were collected for constructing the quaternary structural models, including PDB ID 1FIQ (chains C and A, with a high S-score of 142.8), 3HRD (chains C and D, S-score=98.0), and 1RM6 (chains A and B, S-score=110.9) (Table S6). Functionally, all the three framework templates are related to molybdenum activities, where the 1FIQ is a mammalian xanthine oxidoreductase which parallels yagTSR in its capabilities as an aldehyde oxidoreductase; the 3HRD is characterized as nicotinate dehydrogenase and consists of similar subunits to YagRST, i.e., two larger molybdopterin subunits, one medium FAD-subunit, and a small FeS subunit (34); finally, the 1RM6 is another member of the xanthine oxidase family from *Thaura aromatica*. This enzyme differs however in its enzymatic role, demonstrating affinities towards phenolic compounds rather than aldehydes (28).

The complex structure of YagRST was solved by Correia, et al. (Correia et al. 2016) with a PDB ID: 5G5G, after the structural modeling was performed; this experimental structure can therefore be used as a blind test of the Threpp models. In Figure 5b, we present a superimposition of Threpp-predicted model (in C*α*-trace) and the X-ray structure (cartoon) of the YagRST complex, which has a TM-score=0.90 and interface RMSD=2.01 Å. Here, an interface RMSD was calculated on the C*α* pairs with an inter-chain distance <5 Å, where the Threpp model covers 96.7% of interface residues. This result shows that a close similarity can be achieved between the Threpp model and the native in both global and interface structures.

#### Comparison of Threpp models on 39 solved PPI complexes

In fact, there are in total 39 out of the 35,125 protein-protein complexes whose structures have been experimentally solved in PDB since 2016, which is the time when our PPI structure library was constructed, on which the Threpp structural modeling was based. Compared to these experimental structures, the average TM-score of the Threpp models is 0.73, where the average sequence identity between the target and complex template in our modeling is 48% (see Table S7 for detailed list of the 39 proteins and https://zhanglab.ccmb.med.umich.edu/Threpp/download/solved_structures.zip for PDB format of the structural models). These results further demonstrate the effectiveness of Threpp to the quaternary structure prediction.

## CONCLUSION

We developed a new pipeline, Threpp, for recognizing and structure modeling of protein-protein interactions in organisms. Starting from a pair of monomer sequences, dimeric threading was extended to scan both sequences against a complex structural library collected from the PDB. The alignment score of the dimeric threading was then combined with the high-throughput experimental data through a naïve Bayesian classifier model to predict the likelihood of the target sequences to interact with each other, where the quaternary structure models of the identified PPIs were built by reassembling the monomeric alignments with the quaternary template structural frameworks.

The pipeline was tested on a large set of protein pairs containing 763 experimentally established PPIs and 134,632 non-interacting protein pairs compiled from different cellular compartments. It was shown that although the threading-based assignment does not create a higher accuracy of PPI recognitions than the best high-throughput experiments, the combination of them can result in a significantly higher PPI recognition rate with the Matthews correlation coefficient 18.5% higher than the best dataset from the HTE; this increase is mainly attributed to the complementarity of the threading-based approach to the HTE results.

As an application, Threpp was extended to scan all sequences in the *E. coli* genome and created 35,125 high-confidence PPI predictions, which is 4.5 times higher than that without using the threading-based component scores (7,872). This significant data boost demonstrates the usefulness of complimentary computer-based PPI predictions in the interactome constructions against the high-throughput experiments. A detailed network analysis was performed on the Threpp PPI predictions, which revealed that the degree of the PPI networks follows strictly a power-law distribution. This scale free feature is essential to the robustness of the PPI networks against evolution as the majority of proteins interact only with few partners, which makes the networks less sensitive to the deletion and insertion of local protein nodes. On the other hand, a substantial amount of functionally important proteins, which is significantly higher than that expected from a normal degree distribution, are found in direct interactions with nearly twice more proteins than the non-essential proteins. These proteins serve as the hub of the PPI networks and play an essential role for *E. coli* survival.

To create structure models of the protein-protein interactions, Threpp reassembles the monomer models of each component obtained by single-chain threading approaches on the dimeric framework of the complex from the dimeric threading alignments. 6,771 out of the putative 35,125 PPIs are found to have a high confidence score that corresponds to the correct fold of the complexes with a TM-score >0.5. As a case study, two examples from dimethyl sulfoxide reductase (DmsAB) and trimeric iron-sulfur (YagRST) complexes are examined in detail, where the predicted models are found highly consistent with the experimental data from previous functional studies. Overall, 39 complex structures were solved after the structure library was created, where 72% of them have a TM-score >0.5, resulting in an average TM-score 0.73 compared to the native (Table S7).

Historically, as a major technique of PPI assignments, the HTE has been suffering from high false positive rate at the beginning of its emergence. Meanwhile, traditional structural biology techniques (X-ray and NMR) have more difficulties in determining the PPI complex structures than that encountered for monomer proteins. These difficulties have significantly frustrated the progress of the interactome studies compared to the success of structural genomics that focuses on the structure and function of monomer proteins. The results presented in this study demonstrates promising improvement on both aspects of interactome through a hybrid pipeline that combines computational threading and traditional HTE datasets with analogy-based structure modeling. Although the pipeline has been applied only to *E. coli* in this study, it can be readily extended to the study of other organisms. With continuous improvements of the threading techniques and the enlargement of PPI structure datasets through new techniques such as cryo-EM (Cheng 2015), the Threpp pipeline, which has been made freely downloadable to the community, should find the increasing usefulness on the studies of other interactome systems.

## METHODS

Threpp consists of three consecutive steps of multiple-chain threading, Bayesian classifier-based interaction prediction, and complex structure construction, where the flowchart is depicted in Figure 1.

### Dimer-threading based PPI recognitions

The multi-chain threading procedure in Threpp is extended from a former version of SPRING that was designed to detect complex structure templates for protein pairs of known interactions (Guerler et al. 2013). Initially, one of the target sequences (e.g. Chain A) is threaded by HHsearch, a profile-profile sequence aligner assisted with secondary structure (Wu and Zhang 2008), against the monomeric template library from the PDB, to create a set of putative templates (*T*_*Ai*_, *i*=1,2,…) each associated with a Z-score (*Z*_*Ai*_). Here, the Z-score is defined as the difference between the raw alignment score and the mean in the unit of standard deviation, where a higher Z-score indicates a higher significance and usually corresponds to a better quality of the alignment. In parallel, the opposite chain (e.g. Chain B) is threaded separately by HHsearch through the PDB, yielding a set of templates (*T*_*Bi*_) with Z-score (*Z*_*Bi*_). Then, all binding partners of the *T*_*Ai*_ are gathered from the oligomer entry that is associated with *T*_*Ai*_ in the PDB. If any of the binding partners of *T*_*Ai*_ is homologous to any of the high-ranking templates of Chain B (*T*_*Bi*_), an interaction framework is established for the target complex from the oligomer associated with *T*_*Ai*_ (middle column in Figure 1).

The homology comparisons between the PDB templates are pre-calculated by an all-to-all PSI-BLAST scan where a homology is defined between two templates if the E-value <0.01. The Z-score of the framework is defined as the smaller of the two monomeric Z-scores. For heterodimer proteins, this threading process is repeated using Chain B as the starting probe to identify binding partners and the frameworks. The confidence of the target chain interactions by the threading alignments is evaluated by the highest Z-score of the complex, named *Z*_*com*_, among all the templates identified by the procedure.

### Bayes classifier for multiple evidence combination

To evaluate if the putative chains (A and B) interact, we combined the *Z*_*com*_ score with the interaction evidence from HTEs through a model of the naïve Bayes classifiers (Domingos and Pazzani 1997). With the classifier, Threpp_threading is encoded by a single binary feature which equals ‘1’ if *Z*_*com*_ ≥ 25 and otherwise ‘0’. The threshold of 25 was derived from a separate set of 200 training proteins taken from the PDB, which are non-homologous to the test proteins of this study, by maximizing the Matthew’s correlation coefficient (MCC) of the PPI recognitions. Data from each of the experimental datasets is also represented by a binary feature which equals ‘1’ if the corresponding experiment indicates that the pair of proteins interacts and otherwise ‘0’. In the present study, we identified four experimentally derived PPI networks for *E. coli* from the literature (Butland et al. 2005; Arifuzzaman et al. 2006; Hu et al. 2009; Rajagopala et al. 2014), although more HTE features can be combined similarly when available.

The classifier is parameterized using the positive and negative gold standard sets, which are randomly split into five subsets with equal size, where four of the five subsets are used to estimate the conditional probabilities for the positive (*P*) and negative (*N*) samples by

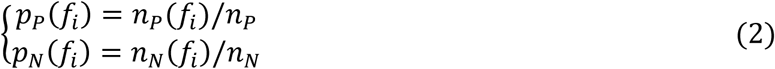

where *n*_*P*(*N*)_(*f*_*i*_) is the number of positive (negative) interacting cases for a given score of *f*_*i*_ of the *i*th feature (*i* = 1, ⋯, 5 represents the five features from Threpp_threading and HTE datasets), and *n*_*P*_ and *n*_*N*_ are the total numbers of positive and negative interacting cases in the training sample, respectively. The remaining subset of protein pairs are used for testing in Results, where the likelihood of interaction is evaluated by

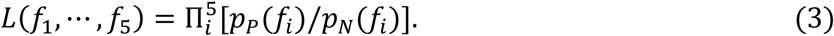

We note that the likelihood ratio is derived solely from features that are available, indicating that the sample protein pairs are interacting. The remaining features are excluded (i.e. treated as missing evidence) since the unavailability of an experimental confirmation or threading alignments does not indicate whether a pair of proteins interacts or not.

### Structure assembly of protein complexes

If the proteins are deemed to interact, the complex structures are constructed by structurally aligning the top-ranked monomer templates of Chain A and B to all putative interacting frameworks using TM-align (Zhang and Skolnick 2005). The structural alignment is built on the subset of interface residues. The resulting models are evaluated by the Threpp score of

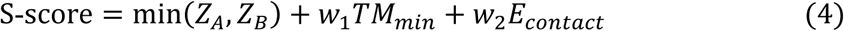

where *Z*_*A*(*B*)_ is the Z-score of the monomer threading alignment by HHsearch for Chain A(B); *TM*_*min*_ is the smaller TM-score returned by TM-align when aligning the top-ranked monomer models of A and B to the interaction framework; *E*_*contact*_ is a residue-specific, atomic contact potential derived from 3,897 non-redundant structure interfaces from the PDB using the formula of RW (Zhang and Zhang 2010). The weight parameters *w*_1_ and *w*_2_ are set to 12.0 and 1.4 through a training set of protein complexes to maximize the modeling accuracy of the interface structures.

## AUTHORS’ CONTRIBUTIONS

Y.Z. conceived and designed the experiments; W.G., A.G. and C.L. performed the experiments and analyzed the data; C.Z. developed the webserver; E.W. helped prepare the figures; W.G., A.G., C.L. and Y.Z. wrote the manuscript.

## AVAILABILITY OF DATA AND MATERIALS

All benchmark and *E coli* PPI modeling data, together with the on-line server of the Threpp method, are made available at PPI modeling data, together with the on-line server of the Threpp method, are made available at https://zhanglab.ccmb.med.umich.edu/Threpp/.

## ETHICS APPROVAL AND CONSENT TO PARTICIPATE

Not applicable.

## CONSENT FOR PUBLICATION

All authors have approved the manuscript for submission.

