## Supporting Information for "Integrating multimeric threading with high-throughput experiments for structural interactome of *Escherichia coli*"

### Supplementary Information

**Tables S1** lists the gold standard PPIs of the *E coli* genome and is available to download at https://zhanglab.ccmb.med.umich.edu/Threpp/download/Table_S1.xlsx.

**Tables S2** lists PPIs predicted by Threpp_threading on the *E coli* and is available to download at https://zhanglab.ccmb.med.umich.edu/Threpp/download/Table_S2.xlsx.

**Tables S3** lists PPIs detected by high-throughput methods on *E coli* and is available to download at https://zhanglab.ccmb.med.umich.edu/Threpp/download/Table_S3.xlsx.

**Tables S4** lists the final PPIs predicted by Threpp on the *E coli* genome and is available to download at https://zhanglab.ccmb.med.umich.edu/Threpp/download/Table_S4.xlsx.

**Tables S5** lists the betweenness centrality for all proteins in *E coli* that have at least one PPI partner in the Threpp predicted networks and is available to download at https://zhanglab.ccmb.med.umich.edu/Threpp/download/Table_S5.xlsx.

**Table S6.** Summary of Threpp modeling parameters for the two PPI complexes shown in Figure 5.

| Target ID | Template ID | Template function | Z-score[a] | S-score[b] |
|---|---|---|---|---|
| *Threading model for monomer chains* | | | | |
| DmsA | 1EU1_A | Oxidoreductase | 56 | |
| DmsB | 2VPZ_B | Oxidoreductase | 32 | |
| YagR | 1RM6_A | CoA reductase (alpha-unit) | 159 | |
| YagS | 1RM6_B | CoA reductase (beta-unit) | 80 | |
| YagT | 3SR6_A | Oxidoreductase | 59 | |
| *Threpp model for complex* | | | | |
| DmsAB | 2IVF_AB | Ethylbenzene/dehydrogenase | 31 | 52.3 |
| YagRS | 1RM6_AB | Oxidoreductase | 79 | 110.9 |
| YagRT | 1FIQ_CA | Oxidoreductase | 58 | 142.8 |
| YagST | 3HRD_CD | Oxidoreductase | 58 | 98.0 |

[a]Z-score for monomer or $Z_{com}$ of Threpp for dimer complex

[b]S-score is defined by Eq. (4)

**Table S7**. Summary of predicted models on 39 proteins solved after the Threpp modeling.

| PDB ID of experimentally solved structures | Comparison of Threpp models to the PDB structures | | | |
|---|---|---|---|---|
| | TM-score | Template ID | S-score | Sequence identity |
| 5CB0_AB | 0.96 | 4R2K_AB | 104.8 | 50% |
| 5T1O_AB | 0.60 | 2XDF_CA | 51.6 | 71% |
| 5VM2_AB | 0.48 | 2DQ4_AB | 106.6 | 49% |
| 5CHN_AB | 0.45 | 1CHM_AB | 128.3 | 48% |
| 4UHT_AB | 0.45 | 1K66_AB | 56.4 | 49% |
| 5JFF_AB | 0.74 | 3ZGY_BA | 42.3 | 77% |
| 6E4B_AB | 0.81 | 3LGA_AB | 80.3 | 48% |
| 5IMJ_AB | 0.91 | 2OEZ_AB | 139.2 | 48% |
| 5HW4_AB | 0.94 | 3KWP_AB | 118.1 | 49% |
| 5DUD_AB | 0.55 | 3GI0_AB | 251.5 | 54% |
| 5DUD_AC | 0.46 | 3GI0_AB | 242 | 46% |
| 5DUD_BD | 0.43 | 3GI0_AB | 198.2 | 47% |
| 6GAM_ST | 0.97 | 4GD3_ST | 122.3 | 43% |
| 6GAM_LM | 0.95 | 4UE3_LM | 124 | 49% |
| 5AEE_AB | 0.43 | 1WE5_AD | 146.8 | 39% |
| 6GFL_AB | 0.49 | 3BQ9_AB | 181.5 | 50% |
| 5WQL_AB | 0.94 | 1XNF_AB | 76.12 | 47% |
| 5Z1Z_AB | 0.87 | 2GO1_AB | 158.5 | 50% |
| 5ZE6_AB | 0.48 | 4JYX_BD | 140.4 | 50% |
| 5NJ9_AB | 0.75 | 1VPB_AB | 139.2 | 47% |
| 5NJ9_BD | 0.49 | 1VPB_AB | 144 | 50% |
| 5XU7_AB | 0.50 | 2WDO_AB | 62.3 | 33% |
| 5J43_AE | 0.96 | 1V7C_AB | 159.5 | 50% |
| 6AGL_AB | 0.49 | 1KFL_AB | 194.1 | 50% |
| 5G5G_AB | 0.90 | 3HRD_CD | 90 | 25% |
| 5G5G_AC | 0.94 | 1FIQ_CA | 142.8 | 28% |
| 5G5G_BC | 1.00 | 1RM6_AB | 110.9 | 31% |
| 6OFU_AB | 0.95 | 1GVF_AC | 136.9 | 50% |
| 6GAM_SL | 0.96 | 4UE3_SL | 243.9 | 50% |
| 4YZE_AC | 0.81 | 3EUP_AB | 115.4 | 49% |
| 5TPM_AB | 0.71 | 2DI3_AB | 111.2 | 45% |
| 6OHB_AC | 0.91 | 3LS9_AB | 124.8 | 48% |
| 5Z03_AB | 0.88 | 3T5M_AB | 146.2 | 48% |
| 6MX1_AB | 0.64 | 3C1N_AB | 214.7 | 49% |
| 6MTG_AB | 0.88 | 2H2W_AB | 113.7 | 52% |
| 5FSR_AB | 0.54 | 2WUQ_AB | 63.3 | 45% |
| 6BPM_CA | 0.67 | 4AIP_AC | 75.9 | 45% |
| 6EI9_AB | 0.41 | 1GWJ_AB | 70.6 | 47% |
| 5ZXL_AB | 0.96 | 1TA9_AB | 118.7 | 49% |
| Average | 0.73 | | 129.4 | 48% |
